# Towards Bayesian-based quantitative adverse outcome pathways using *in vitro* data from open literature and continuous variables: a case example for liver fibrosis

**DOI:** 10.64898/2026.04.15.718674

**Authors:** Robin Durnik, Tereza Juchelkova, Helge Hecht, Levi M.T. Winkelman, Joost B. Beltman, Xavier Coumoul, Florence Jornod, Karine Audouze, Ludek Blaha, Lola Bajard

**Affiliations:** RECETOX, Faculty of Science, Masaryk University, Brno, 611 37, Czech Republic; Division of Cell Systems and Drug Safety, Leiden Academic Centre for Drug Research, Leiden University, Leiden, The Netherlands; Université Paris Cité, INSERM, HealthFex, Paris 75006, France

**Keywords:** hepatic stellate cell activation, collagen, probabilistic model, predictive toxicology, data mining

## Abstract

As toxicology shifts towards non-animal testing, quantitative models are essential to predict adverse health effects from molecular or cellular perturbations. Quantitative Adverse Outcome Pathways (qAOPs) represent such models, building on mechanistic knowledge and quantifying the Key Event Relationships (KERs) described in AOPs. Despite the recognized need, the number of qAOPs remains limited. Bayesian-based approaches are often chosen for developing qAOP for their flexibility, but most use discretized variables, limiting their predictive power. In addition, these models are mainly built from newly generated data, underexploiting the large amount of information available. This study successfully leverages data from public literature and presents an innovative framework based on continuous variables to develop a Bayesian-based quantitative model for a central KER towards liver fibrosis. The model predicts the probability of the expression fold change for two key markers of hepatic stellate cell activation (aSMA and COL1A1), given the effects on tissue injury, using in vitro data from 9 chemicals. We propose a newly developed workflow to assist in knowledge identification, organization, and extraction from scientific literature and chemical databases. Based on *in vitro* data and *in vivo* information from the Open TG-GATEs (Toxicogenomics Project-Genomics Assisted Toxicity Evaluation System) database, we estimate a biologically relevant range in COL1A1 fold change that indicates an activated state of stellate cells and high liver fibrosis odds ratios. Our study provides a case example of integrating published data and continuous variables to build a Bayesian-based model, which constitutes an essential step for predicting liver fibrosis from *in vitro* data.

## Introduction

The need to reduce, refine, and ideally replace animal testing in toxicological studies for ethical, financial, and efficiency reasons has been highlighted for more than a decade^1^, increasing the use of *in vitro* studies based on cellular models. This shift was accompanied by the development of the Adverse Outcome Pathway (AOP) framework.^2^ AOPs provide a structured framework for organizing mechanistic data across different levels of biological organization. They prove valuable at integrating data from *in silico* models, *in vitro* assays and *in vivo* studies, enabling the linking of molecular-level interactions to adverse outcomes (AOs) of regulatory relevance. AOPs begin with a Molecular Initiating Event (MIE) - the interaction between a prototypical stressor (e.g. a chemical) and a molecular target. This triggers a sequence of biological Key Events (KEs), which are connected through Key Event Relationships (KERs). Together, these lead to a final AO relevant to individual or population-level health.^2–4^ Quantitative AOP (qAOP) models are expected to provide a bridge from qualitatively describing knowledge to quantitatively predicting the risk of an AO, which is needed for the regulation of hazardous chemicals.^3,5–8^ A qAOP thus models response–response relationships of all KERs in an AOP, quantitative KERs (qKERs) being the individual pieces.^9^ Although the qAOP concept was introduced about a decade ago^10^, and the importance of qAOPs for risk assessment and regulatory applications has been highlighted in several publications^3,6,9,11^, a limited number have been reported so far (for example, ^5,12–18^). The inherent interdisciplinarity, implying interactions between advanced mathematics and experimental biology, and the availability of appropriate (i.e., properly designed and reported) input data for the models are some of the obstacles restraining qAOP development.

The development of qAOPs can be approached through several strategies. Mechanistic, systems biology–based models allow detailed simulation of molecular and cellular processes, but they are data-intensive, complex, and difficult to generalize.^5^ Probabilistic modeling, particularly Bayesian networks (BNs), offers a more flexible and accessible alternative for qAOP development.^5,9,12,19–22^ BNs are probabilistic graphical models representing causal or associative dependencies among variables in the form of directed acyclic graphs.^23^ Each node in the network corresponds to a variable, such as a KE in an AOP, and edges represent probabilistic dependencies analogous to KERs. BNs are grounded in Bayesian statistics, where prior knowledge about model parameters is updated with observed data using Bayes’ theorem.^24,25^ This framework naturally accommodates uncertainty, missing data, and heterogeneous datasets, which are common in toxicology, and allows probabilistic inference on how perturbations at the molecular or cellular level propagate toward AOs. Most environmental and toxicological BN applications are based on discrete data^26,27^, which simplifies computation but comes with limitations. Discretization introduces information loss, reduces statistical accuracy, and complicates interpretation.^28–30^ In addition, the choice of discretization intervals and cut-offs is difficult to standardize and fully justify on both statistical and biological grounds, but it can substantially affect the results and interpretation of BN models.^31,32^ On the other hand, continuous-variable BNs overcome drawbacks of discretization by modeling graded biological responses and concentration–effect relationships, enabling more accurate probabilistic translations of in vitro measurements into in vivo-relevant predictions, while retaining uncertainty quantification.^30,31,33^ Out of the existing Bayesian qAOPs, only a few use in vitro data and continuous variables,^5,18,34^, limiting their applicability in accompanying the replacement of animal testing and their predictive capacity.

Liver fibrosis was taken as a case example for developing a continuous variable Bayesian-based qKER using in vitro data from published literature. Liver fibrosis is characterized by excessive extracellular matrix (ECM) accumulation in the liver, and advanced liver fibrosis is recognized as a significant risk factor for the development of hepatocellular carcinoma, which is the most prevalent form of liver cancer and ranks as one of the leading causes of cancer-related mortality worldwide.^35,36^ An OECD endorsed (expert-reviewed and validated) AOP exists and forms a solid baseline for qAOP development.^37,38^ The KEs involved in this process are hepatocyte injury/death, tissue resident cell activation, increase of transforming growth factor β (TGFβ), hepatic stellate cell (HSC) activation, and ECM alteration. Computational models (agent-based and Kappa rule-based) simulating molecular and cellular steps leading to liver fibrosis have been published^39,40^, but, to our knowledge, no qAOP focusing on liver fibrosis is available. Here, we present a case example for collecting, curating, and structuring *in vitro* data from the open scientific literature to develop elements of a qAOP for liver fibrosis. We primarily focus our efforts on the KER linking liver cell death, a central KE shared by several AOPs for hepatotoxicity^41^, to HSC activation, an essential step towards fibrosis progression (Figure 1).

**Figure 1:**
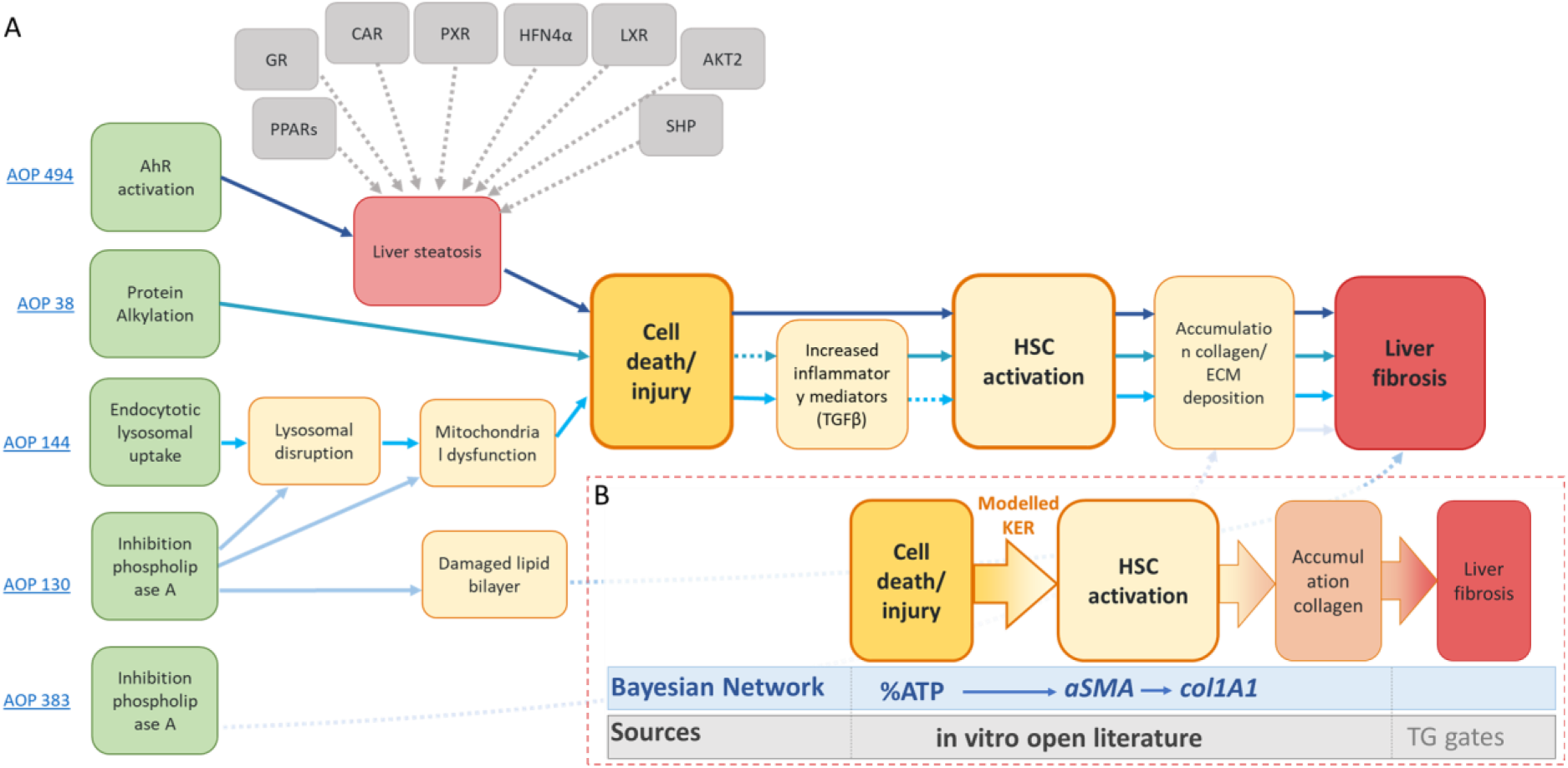
Strategy for developing a central qKER for liver fibrosis. (A) Simplified AOP network for liver fibrosis. Information was obtained from the AOP-wiki (https://aopwiki.org/) as of August 2025, and the simplified version of the AOP network for liver steatosis was derived from Escher and colleagues (2022)^42^. (B) Schematic representation of the central KERs, the BN with the measured variables (with decreased % of ATP indicating higher cell injury, and increased aSMA and COL1A1 indicating higher HSC activation), and the source of data chosen for the quantitative model. Dashed lines indicate indirect KERs (i.e., one or more intermediate KEs are not represented).

## Materials and Methods

### Data collection

The workflow for collecting, sorting, and organizing the data is schematized in Figure 2. Software tools (Publication tracker and Chemical identifier) were developed within this research to facilitate a comprehensive literature search and to prioritize publications based on automatic content screening.

**Figure 2:**
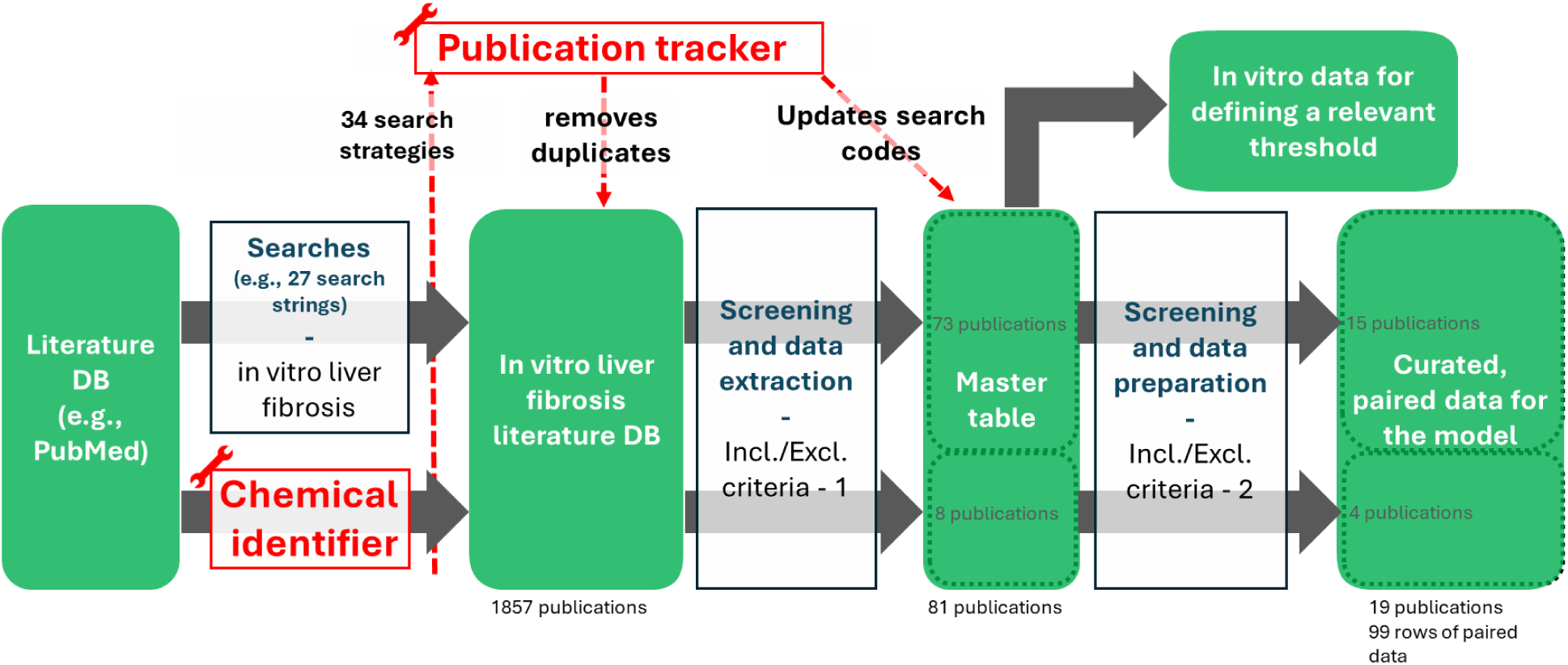
workflow for collecting quantitative data from in vitro literature on chemical-induced liver fibrosis

#### Literature database for chemical-induced liver fibrosis in vitro

Web of Science and PubMed scientific literature databases were screened (date specified in the Appendix 1) to identify publications reporting the effects of chemical exposure on liver fibrosis in vitro using various approaches, using 34 different search strategies.

First, both databases were screened manually using 27 different search strings, most of which retrieved overlapping results (See Appendix 1 for more details). The Publication-Tracker^43^ (Figure 2 and Supplementary Figure S1) tool (using Biopython^44^ package) was developed to systematically track and consolidate results obtained using different search queries, reducing the manual effort required to catalog acquired knowledge.

Second, we developed the Chemical-Identifier^43^ (Figure 2 and Supplementary Figure S2) to prescreen for relevant publications. This tool leverages named-entity-recognition (using Scispacy^45^ and Biopython^44^ Python packages) to identify publications mentioning the studied chemicals compiled for this study from the Tox21^46^ and Open TG-GATEs (Toxicogenomics Project-Genomics Assisted Toxicity Evaluation System)^47,48^. Specifically, 961 studies of liver fibrosis on in vitro multicellular models were retrieved in PubMed using the following search string (optimized using results from the first searches):

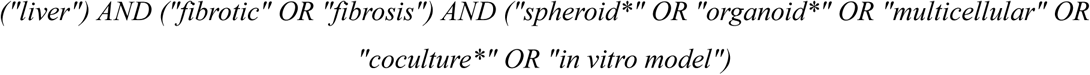

Then, screening with the Chemical-Identifier tool reduced the number of results to 343 unique publications.

Finally, examining publications citing or cited by key publications (see Appendix 1) did not discover new publications, comforting us that the previous searches were comprehensive.

During the process, each search strategy was assigned a search code with a description of the rationale behind it recorded in the search-procedure table (Appendix 1). A database of screened publications (Appendix 2) was established successfully using the Publication-Tracker^43^ tool to avoid duplicate readings and to track the search code(s) that retrieved the given publication.

#### Master table of relevant quantitative data extracted from the literature

Full-text publications from the “chemical-induced in vitro liver fibrosis” database were then filtered using the following inclusion/exclusion criteria: (1) only data normalized to a control group were considered, (2) only data from monocellular *in vitro* models, consisting of stellate cell lines (e.g., LX-2), and from multicellular models, consisting of hepatic cell lines (e.g., HepG2), stellate cells (e.g., LX-2), and in some scenarios, Kupffer cell lines (e.g., THP-1) were included. 81 papers were selected after this filtering step, 73 identified using the manual searches, and an additional 8 found thanks to the Chemical-Identifier. Relevant quantitative data were extracted manually from the selected publication through full-text examination and collected in a master table (Appendix 3). Specifically, for each paper, each exposure condition (given concentration and length of exposure), and each cellular model, we collected values for selected markers of the KEs. The markers were selected based on biological relevance, data abundance, and the robustness of the measurement method. We used the decrease in ATP levels as a marker of cell death and upregulated levels of aSMA and COL1A1 mRNA (measured by reverse transcription quantitative PCR (RT-qPCR), all alternative gene names can be found in the table in Appendix 3) as markers of HSC activation. An increase in ECM, such as collagen, protein concentrations (measured by ELISA or Sirius red staining), was used to represent extracellular matrix alteration, but limited data availability and inconsistent methodologies across studies prevented inclusion of this key event in the model.

#### Preparation of input data for the Bayesian model

The input data for the Bayesian model were selected with the following criteria: (1) only studies using exogenous chemicals such as thioacetamide or carbon tetrachloride were used, excluding studies using, e.g., bile acids, serotonin, or palmitic acid, (2) only studies using liver explants/slices or multicellular models with at least human hepatocytes (e.g., HepaRG, HepG2, primary hepatocytes) and HSCs (e.g., LX-2, hTERT). This is because multicellular models composed of different liver cell types are required to properly recapitulate liver fibrosis events in vitro.^49–51^ In total, 19 papers were selected for the Bayesian model.

### BN elaboration

#### Data preprocessing

Outlier detection followed Tukey’s rule, with values beyond *1.5 × IQR* (interquartile range) considered outliers.^52^ Distributional properties were assessed using Kullback-Leibler divergence (KL), a general measure of divergence between distributions, here applied with a normal distribution as the reference, Q-Q plot correlation, and the Shapiro–Wilk test, providing both quantitative indices and diagnostic plots to inform suitability for probabilistic modeling. All analyses and visualizations were conducted in RStudio (R 4.5.1)^53^ with custom scripts and standard packages (https://github.com/Turmee/qAOP_bayesian_networks).

#### Bayesian network construction

The modeling process consisted of structure definition, parameter learning, and model validation. The BN structure was defined a priori based on established biological knowledge described in AOPs for liver fibrosis, with a particular focus on the endorsed AOP id38^38^, ensuring causally consistent directionality (https://aopwiki.org/aops/38, Figure 1). Specifically, cellular injury (measured by intracellular ATP depletion) was treated as an upstream event influencing stellate cell activation (measured by aSMA and COL1A1 gene expression), representing a central KER linking hepatocyte death to fibrotic response (Figure 1B). A Gaussian Bayesian Network (GBN) was implemented for continuous nodes, where each child node is modeled as a linear function of its parents plus a normally distributed error term^54^ using bnlearn R package (version 5.0.2).^55^

Predictions in GBNs were represented as conditional distributions:

- ATP_perc_: no parents

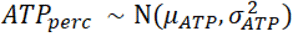
- aSMA: linear regression on ATP_perc

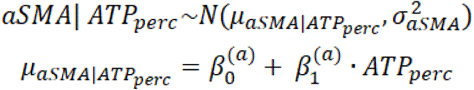
- COL1A1: linear regression on aSMA

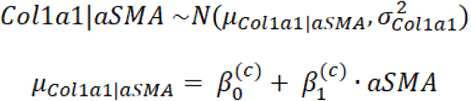

#### Model validation and performance

Performance describes how close model predictions are to observed outcomes, whereas uncertainty describes how concentrated or dispersed the posterior predictions are for a given input. A desirable model combines high predictive performance with low posterior uncertainty. BN performance and uncertainty were assessed using metrics recommended by Marcot and colleagues.^56^ Predictive skill was quantified via posterior predictive correlation (R²) and normalized mean squared error (NMSE), computed under 5-fold cross-validation. Continuous-scale predictions were obtained by likelihood-weighted sampling using the *cpdist()* function from bnlearn package, providing posterior predictive distributions for aSMA and COL1A1. Additional validation included predicted-versus-observed plots to evaluate calibration and residual structure. Predictive uncertainty was characterized by posterior variance and Bayesian credible intervals were used to assess parameter uncertainty.

#### Probabilistic inference

Once the BN was fitted, probabilistic queries were performed to estimate the likelihood of downstream events given evidence on upstream variables. All conditional predictions were generated using cpdist() by sampling from the posterior predictive distribution under specified evidence. For each observation in the cross-validation folds, the sampled distribution of COL1A1 was used to compute the probability of falling into “low” (COL1A1 < 1.5) or “high” (COL1A1 ≥ 1.5) intervals, with the threshold of 1.5 selected based on in vitro and in vivo evidence (detailed in the results section). Quantitative KER (qKER) curves were then obtained by evaluating the BN over a range of ATPperc values and summarizing the corresponding posterior predictive distributions of aSMA and COL1A1, providing a continuous, probabilistic mapping of how perturbations in upstream key events propagate downstream along the modeled pathway. Analogous queries were conducted for aSMA, providing a consistent probabilistic mapping across the modeled pathway, using the threshold of 1.5, as for COL1A1.

#### In vivo data analysis

*In vivo* fold-change data (from Open TG-GATEs) were analyzed to contextualize in vitro predictions and to define relevant thresholds for changes in COL1A1 or aSMA expression potentially associated with higher odds ratios (ORs) of liver fibrosis. Group-wise differences between samples with or without liver fibrosis (as classified in TG-Gates histology reports) were assessed using the Wilcoxon rank-sum test. Logistic regression models were then fitted separately for each marker to evaluate the relationship between HSC activation and fibrosis. Specifically, fibrosis status was modeled as a function of either aSMA fold change (Fibrosis ∼ aSMA) or COL1A1 fold change (Fibrosis ∼ COL1A1), filtering transcriptomics data to include only observations with a significant fold change in aSMA or COL1A1 gene expression (p-value < 0.1), respectively. The magnitude and statistical significance of associations between gene expression and fibrosis were evaluated separately for aSMA and COL1A1. The resulting ORs therefore quantify the change in odds of fibrosis per unit increase in fold change for each marker independently.

Logistic regression was then used to model the odds of fibrosis as a function of HSC activation, with both aSMA and COL1A1 fold change included simultaneously as predictors (Fibrosis ∼ aSMA + COL1A1). ORs therefore represent adjusted effects, quantifying the change in odds of fibrosis per unit increase in fold change while controlling for the other marker. Data were filtered to include only those with a significant fold change in expression (p-value <0.1). Although both aSMA and COL1A1 fold change are used in the model, the magnitude and significance of the associations between gene expression and ORs were examined separately for each gene.

## Results

We chose to focus on the KER linking hepatic cell death to hepatic stellate cell activation, as it has a central role, being shared by several AOPs for liver fibrosis (Figure 1).

### Input data for the Bayesian model

#### Data collection and extraction

Quantitative data were extracted from 19 relevant publications about chemical-induced liver fibrosis in multicellular in vitro models, identified through a structured, comprehensive search assisted by newly developed software tools (see Material and Methods and Figure 2). The Chemical Identifier, in particular, retrieved 17 of the 19 publications, including 4 studies not identified with the classical approaches (either by search string or by being cited by another relevant study). The two missing publications were not detected due to the absence of the chemical name in their abstracts. Overall, the Chemical-Identifier filter narrowed down 343 publications containing relevant chemicals (out of 961 studies of liver fibrosis on in vitro multicellular models retrieved from PubMed), while keeping the majority of the relevant publications. Values reported for markers of each KE were collected and paired per publication and exposure condition (99 individual rows in the master table). The characteristics (e.g., distribution, variable correlation) of these data extracted from the literature were then analyzed before being used to build a Bayesian-based model.

#### Data description

The final dataset comprised *in vitro* measurements obtained from multicellular liver models exposed to nine hepatotoxic chemicals: acetaminophen, methotrexate, thioacetamide, carbon tetrachloride, 2,3,7,8-tetrachlorodibenzo-p-dioxin (TCDD), benzo[a]pyrene, polychlorinated biphenyl 126 (PCB126), allyl alcohol, and ethanol. Chemical (in)dependency, a key feature of AOPs, could not be examined due to a limited amount of data per chemical. Models were tested at multiple concentrations and exposure durations. All three variables, ATP content (ATP_perc_), aSMA expression (aSMA), and COL1A1 expression (COL1A1) were treated as continuous; their descriptive statistics (mean, standard deviation (SD), min, max) are shown in Supplementary Table S1.

In total, 28 complete cases were retrieved with all three primary endpoints measured simultaneously (from 99 recorded rows), while partial data for individual endpoint pairs increased the total usable sample size for modeling of individual steps (see Table 1 for completeness summary). A total of 14 outliers were identified and excluded from further analysis (boxplots of all data can be found in Supplementary Figure S3).

**Table 1.**
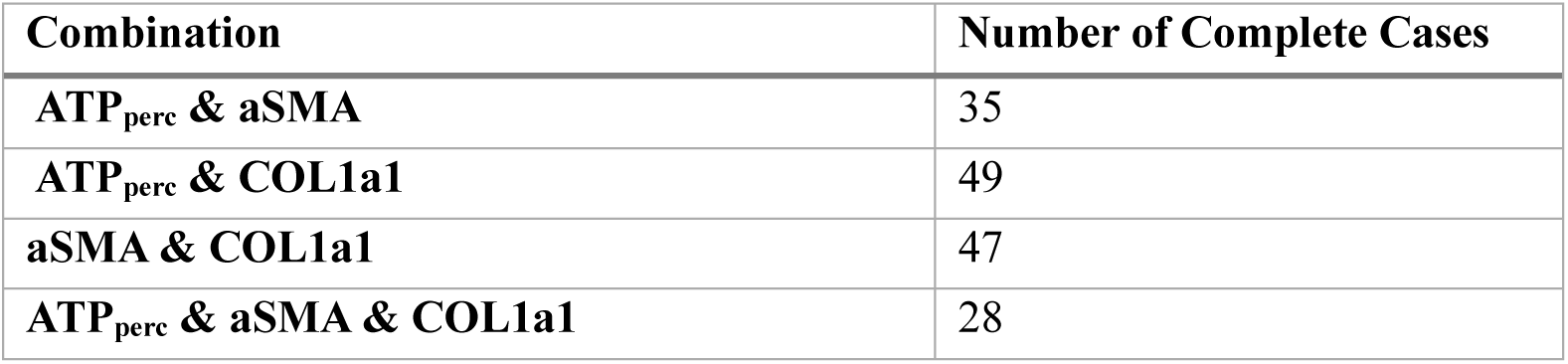
Summary of data completeness for selected variables.

| Combination | Number of Complete Cases |
| --- | --- |
| ATP<sub>perc</sub> & aSMA | 35 |
| ATP<sub>perc</sub> & COL1a1 | 49 |
| aSMA & COL1a1 | 47 |
| ATP<sub>perc</sub> & aSMA & COL1a1 | 28 |

ATP levels were quantified as ATP_perc_ and used directly in all statistical analyses and model fitting. For visualization and interpretative purposes only, ATP values were re-expressed as a measure of cell injury, defined as max(ATP_perc_) − ATP_perc_, such that higher values correspond to increased cellular damage.

#### Distributional characteristics

All three biological endpoints showed deviations from perfect normality, consistent with the expected heterogeneity of *in vitro* responses (Supplementary Figure S4). ATP_perc_ exhibited an approximately symmetric distribution with slightly heavy tails (KL > 0.2), while aSMA and COL1A1 displayed stronger skewness (KL divergence > 0.2). Shapiro–Wilk tests returned p < 0.05 for all endpoints, indicating statistically significant deviations from normal distributions. No transformations were applied, as these did not improve normality (data not shown) and would have complicated interpretation.

#### Correlations and variable relationships

Pairwise correlations among the three primary endpoints were examined using Pearson correlation. Uncertainty was quantified using nonparametric bootstrap resampling (10,000 replicates), in which observations were sampled with replacement and correlations recomputed for each replicate. Confidence intervals were derived using the percentile method (Figure 3). Across all data, cell injury showed a positive association with aSMA (r ≈ 0.554 95 % CI (Confidence Interval) [0.21, 0.78]) and with COL1A1 (r ≈ 0.57, 95 % CI [0.33, 0.78]), consistent with the biological expectation that cell injury is associated with fibrotic activation. The correlation between aSMA and COL1A1 was positive (r ≈ 0.70, 95 % CI [0.31, 0.90]). This is consistent with both genes being markers of HSC activation.

**Figure 3:**
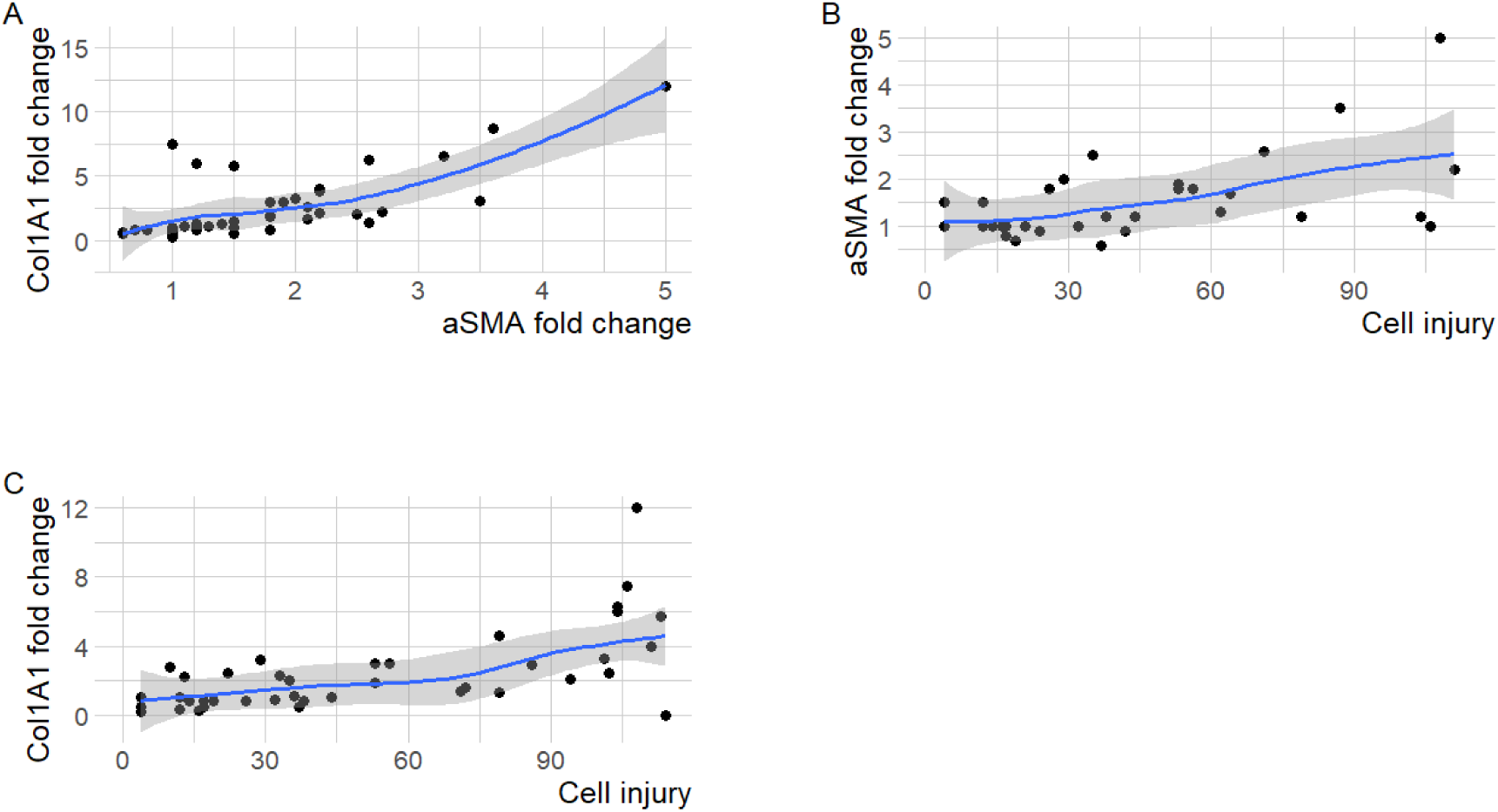
Variables correlations. Scatter plot of variable pairs (fold change of aSMA and COL1A1, cell injury (max(ATP_perc_) − ATP_perc_)) showing individual observations, with a LOESS (Locally Estimated Scatterplot Smoothing) curve (blue) indicating the overall trend and grey shading representing the 95 % confidence interval of the estimates.

The influence of exposure duration was examined as a grouping factor using Multivariate Analysis of Variance (MANOVA) to identify the optimal threshold maximizing separation in aSMA and COL1A1 responses in both in vivo and in vitro data. However, the substantial imbalance in group sizes, with a limited number of observations in some subgroups, restricts the robustness of these results (Supplementary Figure S5A, B). In addition, the relationship between exposure duration and aSMA or COL1A1 expression was explored using regression-based and non-parametric approaches. Due to the sparsity of the data and high variability, no consistent or statistically supported trend could be established across the tested models (Supplementary Figure S5C, D).

The data extracted, collected, and analyzed were then used to build the Bayesian model.

### Bayesian model for the qKER “hepatic cell death leading to stellate cell activation”

We constructed a GBN model to represent causal relationships among the three KEs. The final structure (ATP_perc_ → aSMA → COL1A1) reflected established mechanistic understanding of hepatocellular injury leading to stellate cell activation (Figure 1B). Although both aSMA and COL1A1 increased expression are generally used as markers of HSC activation, we placed COL1A1 downstream of aSMA for two reasons. First, one in vivo study in rodents suggested that aSMA may activate COL1A1.^57^ Second, COL1A1 potentially represents a transition to the consecutive KE that largely consists of collagen protein accumulation.

Based on standard recommendations for power analysis, a minimum of 20–30 observations are required for the proposed three-node model, while 50–100 observations are ideal for stable parameter estimation, placing this dataset (with 35 to 49 observations, see Table 1) close to the lower, yet acceptable limit. Model parameters were estimated via maximum likelihood using the *bnlearn* package, with each continuous node modeled as a linear regression on its parent(s) and a Gaussian residual term. (Supplementary Table S2 and Supplementary Figure S6). The regression of aSMA on ATP_perc_ shows a negative association with an estimated coefficient 𝛽_1_^(𝑎)^ = − 0.015 and a 95 % credible interval ranging from -0.030 to -0.004. This confirms that lower ATP levels (higher cell injury) are associated with higher aSMA expression (higher HSC activation), and the narrow credible interval indicates this relationship is relatively stable and well-supported by the data. For COL1A1 on aSMA 𝛽_1_^(𝑐)^ = 1.941 (95 % CI [0.85, 2.60]), indicating that increased aSMA levels predict increased COL1A1 expression. The effect is supported as the credible interval excludes zero.

Predictive performance was quantified using NMSE and R², with uncertainty estimated from 200 bootstrap resamples (Table 2). For aSMA, the model explained very little variance (R² = 0.13, NMSE = 0.93), indicating that ATP_perc_ captures only limited variability in aSMA expression. In contrast, COL1a1 predictions were more accurate (R² = 0.31, NMSE = 0.74), consistent with the strong correlation between aSMA and COL1a1 (r = 0.69).

**Table 2:** Performance metrics for the aSMA and COL1A1 nodes estimated from 200 bootstrap fits of the BN model, reported as mean ± SD for NMSE, and R².

| Parameter | NMSE | R2 |
| --- | --- | --- |
| aSMA | 0.93± 0.21 | 0.13± 0.13 |
| COL1A1 | 0.74± 0.17 | 0.31± 0.19 |

For thresholded outcomes (aSMA_high ≥ 1.5, COL1a1_high ≥ 1.5), the Bayesian network showed modest to moderate discrimination ability: the area under the ROC curve (AUC) was 0.664 for high aSMA and 0.750 for high COL1A1.

These results align with expectations for small-sample biological datasets, where even moderate R² values (≈15–30 %) are considered meaningful.^58^ Calibration plots comparing predicted versus observed values showed generally linear alignment, with no major systematic bias across the prediction range (Supplementary Figure S7).

Overall, the network captured biologically plausible dependencies, supporting the proposed causal chain in which hepatic cell injury, measured by ATP depletion, leads to aSMA activation, which in turn promotes COL1A1 expression, consistent with the AOP network described in Figure 1. In the following steps, we put more emphasis on COL1A1 as a marker of hepatic stellate activation, but also as a gene encoding an ECM component, putting it more downstream towards the AO.

The Bayesian-based model built using the in vitro data collected from the literature was then used to predict aSMA and COL1A1 fold change given the level of cell injury.

### Biologically relevant threshold of COL1A1 gene expression fold change

As the next step, we aimed to quantify how upstream perturbations propagate along the pathway, by performing conditional probability queries. This allows for calculating the probability of one variable (e.g., COL1A1 mRNA expression) being within a certain range, given the value of the conditioning variable (e.g., % of ATP). To define a biologically relevant range for the fold change in COL1A1 expression, we used two approaches, i.e. approaches to define thresholds for COL1A1 expression associated with *in vitro* HSC activation, and with *in vivo* fibrosis association.

#### Defining a threshold of COL1A1 expression fold change indicative of HSC activation

First, to define COL1A1 expression levels indicative of an activated state, we used *in vitro* studies that displayed stellate cell activation in multicellular cultures or stellate cell monocultures (mostly LX-2) triggered by TGFβ. These studies did not use other chemical exposures and often intentionally aimed to induce stellate cell activation (e.g., to establish experimental models of liver fibrosis).

Values for COL1A1 expression fold change in hepatic stellate cells (LX-2 cell line) after exposure to various levels of TGFβ, as reported in the open literature, were plotted (Figure 4A). There was no clear dose-response, possibly since the response quickly saturates, i.e., all concentrations used are causing maximum activation. In addition, the data were also quite spread, likely related to heterogeneity in experimental conditions, with few studies reporting weak or no fold change, limiting their usefulness for defining a relevant threshold. The latter could be due, at least in part, to spontaneous activation of stellate cells when cultured in vitro.^50,59^ Indeed, if stellate cells are already in an activated state, exposure to TGFβ would not induce a strong activation, and differences in spontaneous activation among different studies could explain the variability in the gene expression. However, despite the variability, for the vast majority of data points (54 of 59), COL1A1 expression fold change is above 1.5 (Figure 4A). This suggests that a fold change in COL1A1 expression above this threshold might indicate an HSC’s activated state, as also supported by a recent study.^60^ Similar results were obtained for aSMA (Supplementary Figure S8A).

**Figure 4:**
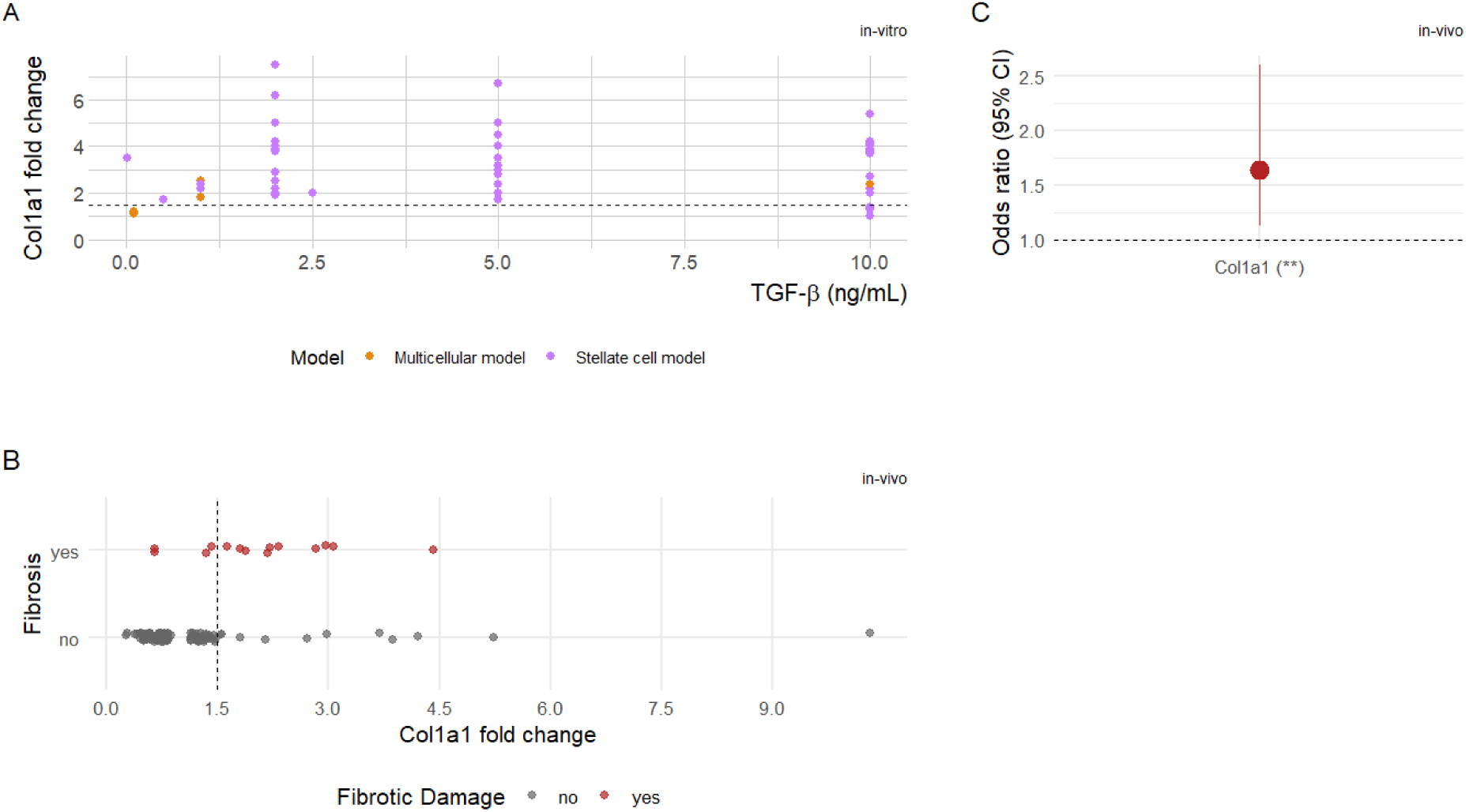
Evaluating biologically relevant thresholds for COL1A1 increase. A) Fold change in COL1A1 expression as a function of levels of TGFβ used to activate stellate cells in vitro, in mono and multicellular *in vitro* models. One data point corresponds to a value reported in one specific study for a given concentration. B) Fold change values of COL1A1 from TG-Gates in vivo data plotted against damage status (Fibrosis), with points for individual observations. Points are colored by damage presence (grey = no, red = yes). C) Odds ratio with 95 % confidence intervals from logistic regression models for COL1A1 fold change in vivo. Asterisks indicate statistical significance (p < 0.05) in neither (-), one (*) or both (**) tests (see Supplementary Table S4). Dashed lines in A and B indicate the 1.5 fold change threshold.

#### Defining a threshold of aSMA and COL1A1 expression fold change indicative of liver fibrosis - integration with in vivo data

Second, to define a threshold in COL1A1 gene expression fold change that would associate with OR of liver fibrosis *in vivo*, data for fold-change gene expression were obtained from the Open TG-GATEs *in vivo* dataset. The dataset contained 3,528 observations from 160 experiments involving various chemicals, concentrations, exposure durations, and exposure types (single vs. repeated), and four types of necrotic damage, one of which was fibrosis. The single- and repeated-exposure groups were well balanced in sample size, although fibrosis cases with significant fold change in COL1A1 or aSMA expression were relatively few. This is because fibrosis information was missing for most samples, with a minority of positive cases, of which only a subset showed a significant fold change in gene expression. After filtering for marker significance (p < 0.1), 14 samples showed significant COL1A1 expression, 9 samples showed significant aSMA expression, and 8 samples were significant for both markers (more information on the filtered dataset can be found in Table S3). Plots of in vivo gene expression fold change based on the fibrosis status indicate that fibrosis cases tend to have increased COL1A1, but not aSMA expression (Figure 4B and Supplementary Figure S8B, C).

Logistic regression and Wilcoxon tests were performed separately for aSMA and COL1A1. For aSMA, both analyses indicated a weak, non-significant, negative association with fibrosis in pooled data (Supplementary Figure S8D and Table S4). On the other hand, COL1A1 exhibited a strong, consistent, and significant positive association in both tests (Figure 4C and Supplementary Table S4), with a mean of 2.1± 1 for COL1A1 fold change in cases with liver fibrosis. To identify a data-driven threshold separating fibrosis vs. non-fibrosis, we focus on COL1A1 as it showed a significant association with the AO, and for which we can therefore define a biologically relevant threshold. We applied a receiver operating characteristic (ROC) analysis using Youden’s J, a criterion that maximizes the combined sensitivity and specificity^61,62^. A COL1A1 fold change threshold of 1.32 yielded the highest performance (J = 0.59, indicating moderate overall discriminatory performance).

In conclusion, analysis of both in vitro data from the open literature and in vivo data from TG-Gates indicates that COL1A1 expression fold changes of 1.5 and above are associated with higher HSC activation (in vitro) and liver fibrosis ORs (in vivo).

### Probabilistic inference of COL1A1 expression using the Bayesian model

Based on these previous results for COL1A1 from in vivo and in vitro TGFβ-induced activated stellate cells, a threshold of 1.5-fold change indicative of HSC activation and higher fibrosis ORs was adopted for the probabilistic inference in the BN. It aligns in vitro and in vivo response ranges and provides a practical boundary between “low” and “high” expression levels. The probability of COL1A1 change in expression being low (COL1A1 < 1.5) or high (1.5 ≤ COL1A1) was computed as a function of ATP_perc_. The resulting probability curve revealed a monotonic decline and increase in the likelihood of low and high COL1A1, respectively, indicating a higher chance of stellate activation and progression towards liver fibrosis, as cell injury increased (Figure 5). Analogous inference for aSMA given ATP_perc_ yielded a similar relationship (Supplementary Figure S9). These quantitative response curves represent qKERs within the AOP framework, translating continuous mechanistic responses into interpretable conditional probabilities.

**Figure 5:**
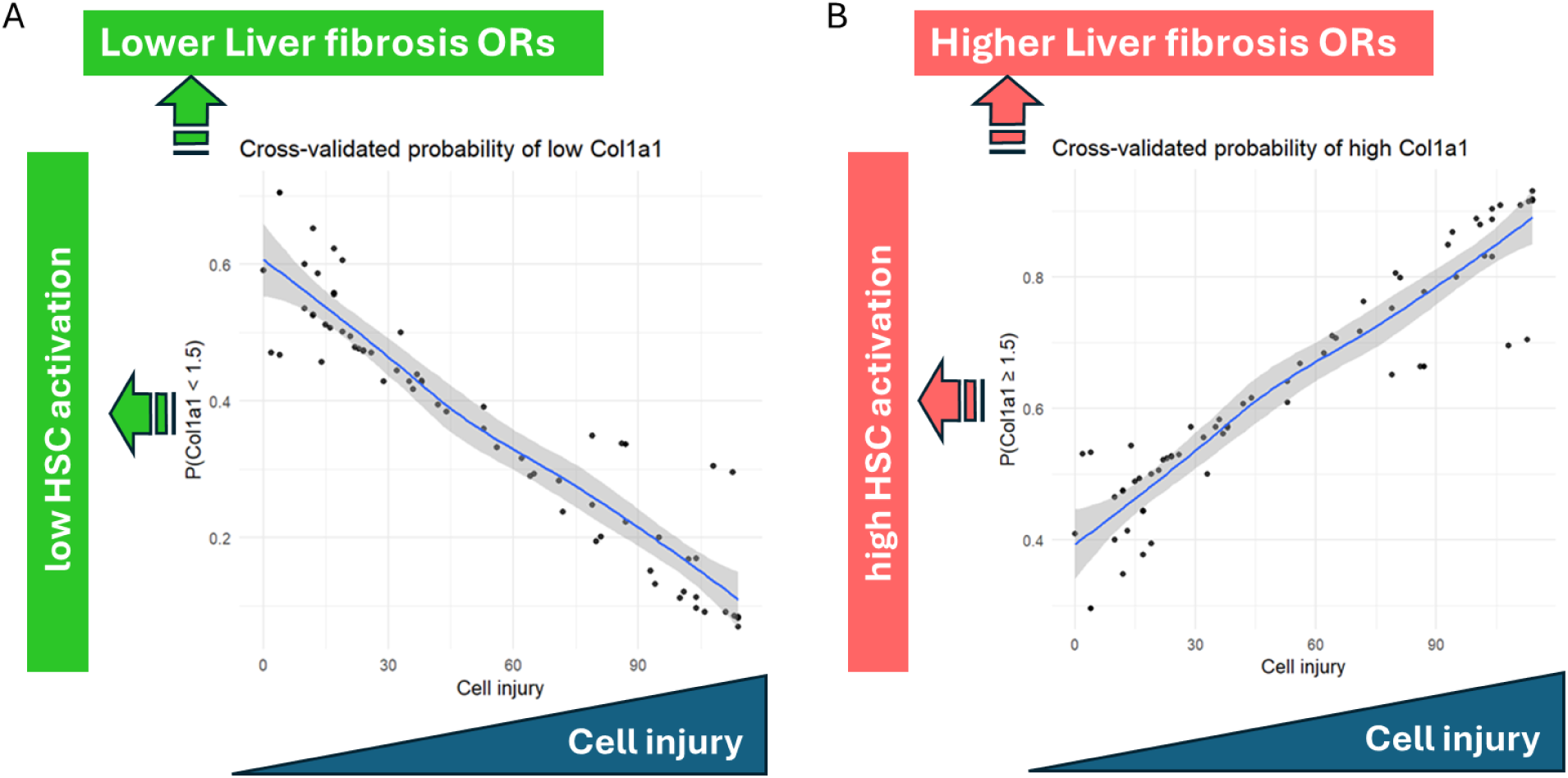
Conditional probabilities of COL1A1 levels given cell injury (max(ATP_perc_) − ATP_perc_) estimated from the Bayesian network model. A) COL1A1 < 1.5 (lower HSC activation/liver fibrosis risk) and B) COL1A1 ≥ 1.5 (higher HSC activation/liver fibrosis risk). Probabilities were obtained using 5-fold cross-validation. Points represent individual cross-validated predictions, blue curves show LOESS smoothing, and grey shading indicates the 95% confidence interval across folds.

### Model interpretation and uncertainty

Predictive uncertainty was quantified using likelihood-weighted sampling, allowing generation of posterior predictive distributions for each node. Model sensitivity analysis—perturbing each variable in turn and monitoring downstream responses—confirmed that ATP_perc_ exerted the stronger and less uncertain influence on aSMA than on the COL1A1, reflecting the upstream position of aSMA in the causal pathway. The aSMA to COL1A1 relationship showed a steeper slope, consistent with aSMA acting as a mediator of ATP-dependent fibrotic activation. Uncertainty bands widened along the chain ATP_perc_ → aSMA → COL1A1 and Bayesian credible intervals around parameter estimates were narrow for the aSMA | ATP_perc_ link but wide for COL1A1 | aSMA, demonstrating accumulation of predictive uncertainty in downstream nodes.

### Limitations

Although we opted for our literature search to be as comprehensive as possible, some relevant publications could have been missed, including those published after the date of the search (as specified for each search in Appendix 2). For instance, two studies reporting the effect of TGFβ on COL1A1 expression in LX-2 were not identified and included in the study,^60,63^ but they support our conclusion on the biologically relevant threshold for COL1A1 expression associated with HSC activation.

Our quantitative model represents a crucial and central KER, but pieces could be added to fully capture the biological complexity underlying liver fibrosis induction and progression, such as the alteration of the ECM. The present study was limited to the most frequently utilized measurements of the KEs (ATP levels, aSMA, and COL1A1 gene expression) to leverage a sufficiently large number of studies using the same methodology, but additional markers might provide a more accurate assessment of the biological effects, in particular, considering the variability in aSMA and COL1A1 gene expression fold changes.

Furthermore, the data were too limited to properly examine the effect of exposure duration. For a chronic disease, such as liver fibrosis, it will be important to integrate time aspects into the model using, for example, ordinary differential equations or Dynamic Bayesian Networks, as proposed in previous studies using either experimental or simulated data.^5,64^ These approaches are also well-suited for incorporating dynamic behaviors of KEs over time, including feedback loops, such as the positive one by which TGFβ signaling increases its own signaling, which might also refine the model.^8,65,66^

## Discussion

We successfully developed a qKER shared by several AOPs for liver fibrosis, providing Bayesian-based predictions of the probability of stellate cell activation (fold-change in COL1A1 expression above a certain threshold) given cell injury (as measured by the percentage of ATP). This novel information was implemented into the AOP-wiki, by updating the pages of the KERid68 (“Cell injury/death leads to Activation, Stellate cells”) and the AOPid494 (“AhR activation leading to liver fibrosis”). Other AOPs that include this KER could not be directly edited at AOP-wiki because we are not co-authors. Using in vitro data from TGFβ-activated stellate cells and in vivo data from Open TG-GATEs, we estimated a biologically relevant threshold of 1.5 for gene expression fold change that would be indicative of stellate cell activation and a higher OR of liver fibrosis. However, thresholds can be adjusted as needed if additional data warrants it. Previous Bayesian-based probabilistic models for liver toxicity have been developed, one specifically for liver steatosis, based on a complex AOP network and using discrete variables, and the other predicting DILI categories based on several in vitro assays performed for the study ^67,19^. Our study is, to our knowledge, the first describing an AOP-anchored, Bayesian-based probabilistic model for liver fibrosis using continuous in vitro data collected from open literature. Our model predictions cannot be directly compared with the previously mentioned, as they differ in their input variables and predicted outcomes.

With the extensive and ever-increasing information published in the scientific literature, text mining tools are needed to ease and accelerate data extraction, for example, for qAOP development.^68–70^ For this study, we developed two complementary software tools - Publication-Tracker^43^ and Chemical-Identifier^43^. Publication-Tracker was used to systematically track publications, while Chemical-Identifier proved useful in reducing the time required to identify key studies by filtering publications based on the presence of relevant chemicals. This newly developed function addresses a limitation of literature search platforms such as PubMed and Europe PMC, where the restricted number of characters and expressiveness of search terms makes it impossible to include extensive lists of chemical names. Beyond the use case presented here, Chemical-Identifier can also be used to explore chemicals associated with a given search term, further supporting comprehensive and efficient literature exploration within the broader toxicology field and beyond.

Models based on continuous variables offer more precise, refined predictions and limit the information loss associated with discretization. Mechanistic models best address these issues, providing deep insight, but their high data demands and complexity limit practical applicability. For example, fully mechanistic qAOPs may involve dozens of differential equations and hundreds of parameters, requiring extensive datasets that are rarely available in practice.^5^ On the other hand, Bayesian-based models offer higher flexibility and better accommodate low data availability, but they mostly use discrete variables. This likely relates to the historical predominance of software optimized for discrete variables, as continuous nodes are either unsupported or require strong assumptions, such as normality or linearity, which may not hold true for experimental data.^21,27^ Even studies claiming to implement continuous or hybrid BNs often discretize data at some stage, either during preprocessing or within the software itself.^22^ Here, we describe a Bayesian-based model using continuous variables. As could be expected, normality and linearity assumptions were not fully met for the available experimental data. To overcome these issues, the model smooths over nonlinear relationships, assumes constant residual variance, and propagates uncertainty linearly through the network. Despite the limitations, the model consistently captures the dominant effects and the accumulation of uncertainty along the ATP_perc → aSMA → COL1A1 pathway, providing relevant predictions. Relationships among nodes in BN are expressed as conditional probability functions—either conditional probability tables (CPTs) for categorical variables or conditional probability distributions (CPDs) for continuous variables.^71^ This flexibility allows integration of many types of data (e.g., in vitro and in vivo data), capturing a wide spectrum of biological variability from mechanistic assays to apical outcomes. A unique property of BNs is their bidirectional inference capability: models can propagate information from causes to effects (parent → child nodes) or from effects to potential causes (child → parent nodes). Combined with their ability to quantify uncertainty, this makes continuous-variable BNs a particularly powerful and easily interpretable framework for qAOP modeling.

Although the extensive work and the use case presented here provides a quantitative model for one central KER, we used in vivo data from Open TG-GATEs to estimate relevant thresholds of COL1A1 gene expression that would associate with a higher OR of liver fibrosis in vivo. With the increasing use of transcriptomics for hazard assessment, identifying biologically relevant thresholds of gene expression fold change associated with functional perturbations becomes particularly important. In addition, several of the chemicals identified in our search as inducing the activation of stellate cells in vitro, are known to induce liver fibrosis in vivo. Thioacetamide, carbon tetrachloride, and allyl alcohol induced liver fibrosis in rat models, as also concluded from the TG-GATEs study. Interestingly, neither ethanol nor acetaminophen led to liver fibrosis in TG-GATEs.^47^ However, they were shown to induce liver fibrosis in other studies – acetaminophen in mice,^72,73^ and ethanol in rats.^74,75^ Compounds such as 2,3,7,8-tetrachlorodibenzo-p-dioxin, benzo[a]pyrene, and methotrexate were not studied in TG-GATEs, but were found to cause liver fibrosis in rat or mouse models in other studies.^76–79^ Similarly, benzo[a]pyrene caused liver fibrosis in mice.^80^ The only negative compound (which did not lead to gene upregulation) in vitro^49^, polychlorinated biphenyl 126 (PCB 126), caused liver fibrosis in mice.^81,82^ The discrepancy could, for example, be related to toxicokinetics or may stem from species differences between humans and rodents. However, the *in vitro* results rely on a single study, which limits the conclusions.^49^ Overall, these observations suggest that stellate cell activation in multicellular in vitro models, as characterized by changes in aSMA and COL1A1 gene expression, and predicted from induced cell injury, may identify chemicals inducing liver fibrosis in vivo. However, evaluating the reliability of in vivo quantitative predictions based on such in vitro model would require additional and better standardized data on a larger set of chemicals.

## Conclusion

Herein, we have established a pipeline for data collection and management for the development of qAOPs. We successfully utilized the newly developed text mining tools (Publication-Tracker^43^ and Chemical-Identifier^43^) that eased data collection and compilation from extensive data sources. To facilitate use by the scientific community, both tools are made accessible via the Galaxy platform (links provided in Supplements).

We demonstrate a case of using in vitro data available in scientific literature to build a quantitative model using Bayesian statistics and continuous variables. Using the information about the extent of cell injury, the model predicts the probability of change in the expression of key markers of stellate cell activation. Using both in vitro and in vivo data, we also derived thresholds in increased gene expression associated with stellate cell activation and liver fibrosis. In addition, the model is anchored on an AOP network, including an OECD-endorsed AOP supported by solid evidence linking downstream key events to the adverse outcome. In the future, the model might be further improved with additional data for a broader range of positive and negative chemicals, experiments providing paired in vitro and in vivo data, as well as quantitative in vitro in vivo extrapolation.

Given the growing importance of FAIR (Findable, Accessible, Interoperable, and Reusable) data and Open Science in today’s research landscape,^83–86^ this work also serves as a case study demonstrating the potential for reusing data from existing scientific literature to make quantitative predictions of downstream in vivo adverse outcomes. It also highlights the usefulness of designing experiments in alignment with AOPs, providing measurements of paired KEs in a given setup, harmonizing procedures (e.g., exposure duration, models, and markers), and transparently reporting to increase the usability for qAOP development.

## Abbreviations

AO, Adverse Outcome; AOP, Adverse Outcome Pathway; MIE, Molecular Initiating Event; KE, Key Event; KER, KE Relationship; qAOP, quantitative AOP; BN, Bayesian Network; TGFβ, Transforming Growth Factor β; HSC, hepatic stellate cell; aSMA, α smooth muscle actin; COL1A1, collagen type I alpha 1 chain; KL, Kullback-Leibler divergence; GBN, Gaussian BN; NMSE, normalized mean squared error; OR, odds ratio; TG-GATEs, Toxicogenomics Project-Genomics Assisted Toxicity Evaluation System

