## Supplementary for "Towards Bayesian-based quantitative adverse outcome pathways using *in vitro* data from open literature and continuous variables: a case example for liver fibrosis"

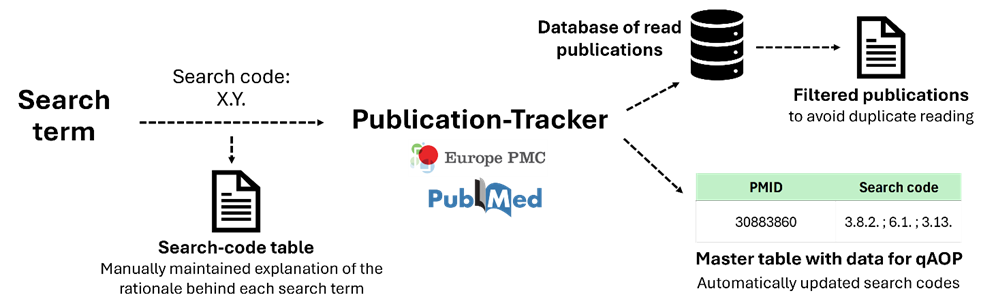

**Supplementary Figure S1** Illustration of the semi-automatized process of publication tracking. Search is conducted in PubMed or Europe PMC via Publication-Tracker that filters for publications you have not yet read. Each search is assigned a search code which links the reasoning behind the search in Search-code table (maintained manually by the user). Search codes are used to track literature in Master table where data used for qAOP development are kept.

<https://usegalaxy.eu/?tool_id=toolshed.g2.bx.psu.edu%2Frepos%2Frecetox%2Faoptk_publication_tracker%2Faoptk_publication_tracker%2F0.1.6%2Bgalaxy0&version=latest>

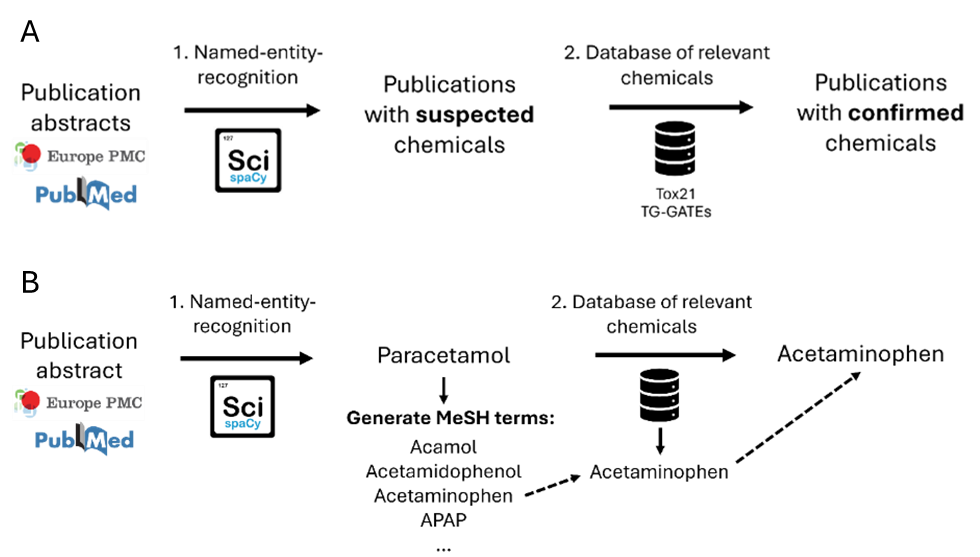

**Supplementary Figure S2:** A) Chemical-Identifier enables identifying literature with toxicologically relevant chemicals. Using this tool, PubMed was screened using large database of chemicals composed of chemicals used in Tox21 and TG-GATEs, reducing the time to find relevant publications. B) An example of how the Chemical-Identifier handles chemical synonyms. If a user-defined chemical database only includes “acetaminophen”, the tool may fail to identify relevant articles that mention “paracetamol”. Using ScispaCy, Chemical-Identifier tries to generate associated MeSH terms - one of which is “acetaminophen”. Since this MeSH term matches an entry in the user-defined database, the publication is correctly flagged as relevant, even though the original text used a different synonym.

<https://usegalaxy.eu/?tool_id=toolshed.g2.bx.psu.edu%2Frepos%2Frecetox%2Faoptk_chemical_identifier%2Faoptk_chemical_identifier%2F0.1.6%2Bgalaxy0&version=latest>

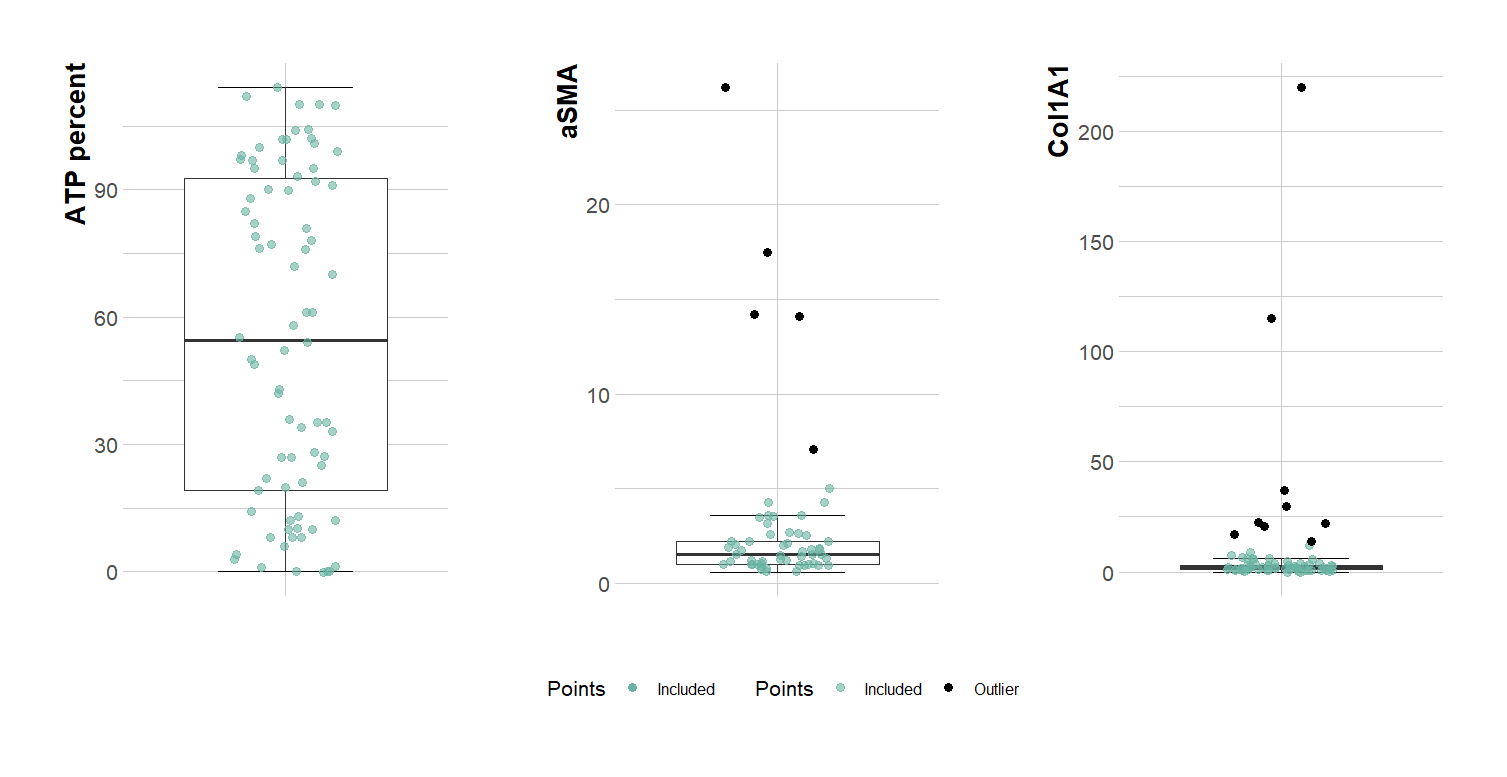
**Supplementary Figure S3:** Boxplots showing distribution of ATP percent, aSMA, and Col1A1 values used for the qKER development. Black points indicate identified outliers. Whiskers indicate variability outside the upper and lower quartiles.

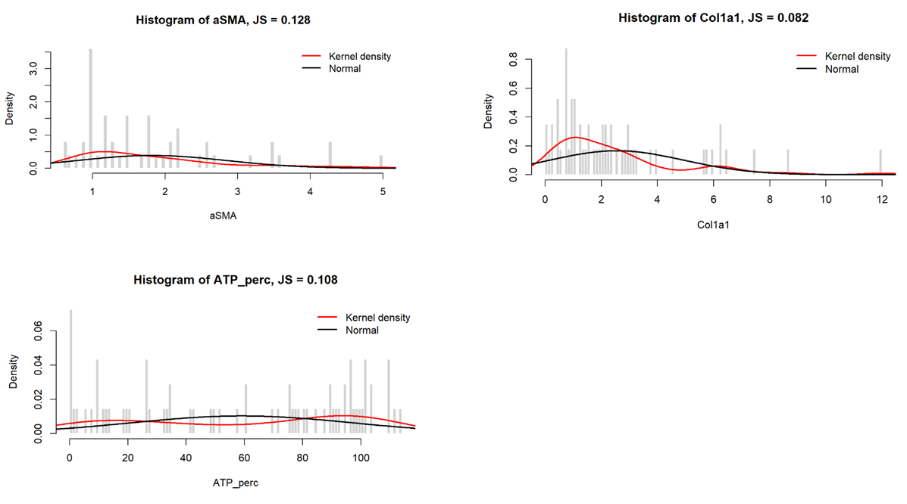

**Supplementary Figure S4:** Histograms of selected variables with kernel density (red), estimated directly from the observed data, and normal density (black), based on the sample mean and standard deviation. Jensen–Shannon divergence between the densities is indicated.

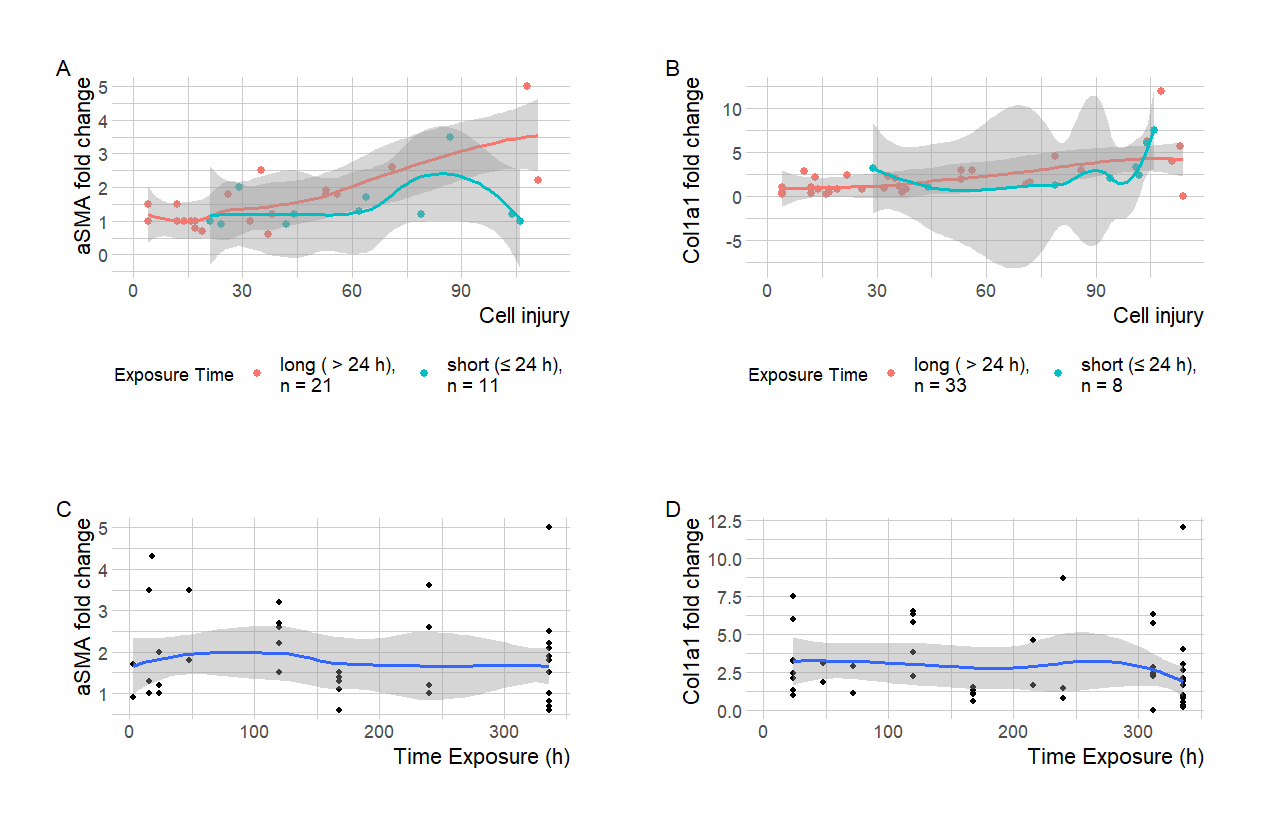

**Supplementary Figure S5: Relationships between cell injury, exposure duration, and fibrotic markers.**
(A–B) Scatter plots show individual observations of aSMA (A) and COL1A1 (B) fold changes versus cell injury, colored by exposure duration (in green, “short” ≤ 24 h; in red, “long” > 24 h). LOESS curves indicate group-specific trends, with shaded areas representing 95% confidence intervals.
(C–D) Scatter plots of exposure time versus aSMA (C) and COL1A1 (D) fold changes, with LOESS smoothing. LOESS curves (in blue) indicate trends, with shaded areas representing 95% confidence intervals.

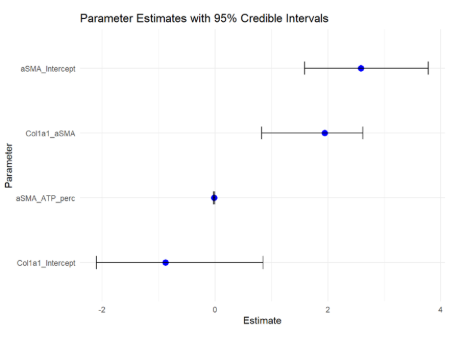

**Supplementary Figure S6:** Visualization of bootstrap-based parameter estimates for the Bayesian network model, shown as point estimates with horizontal uncertainty bars corresponding to 95 % credible intervals.

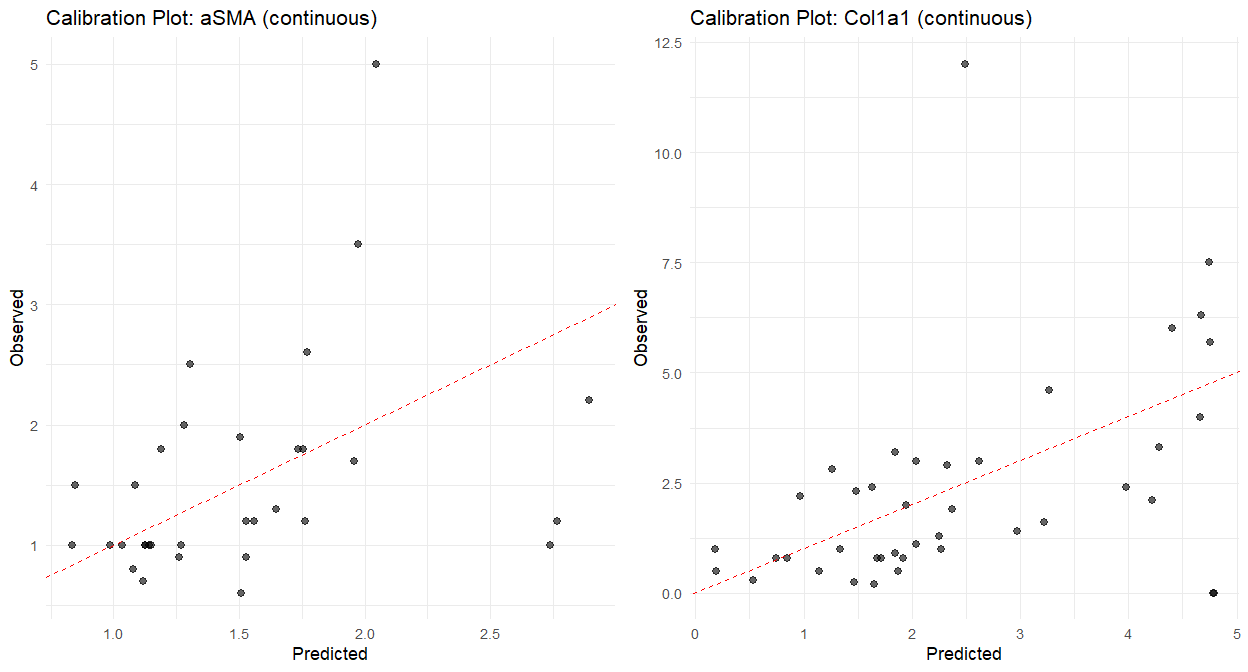

**Supplementary Figure S7**: Calibration plots for the aSMA and COL1A1 nodes in the Bayesian network model, showing predicted versus observed values with a reference line indicating perfect calibration.

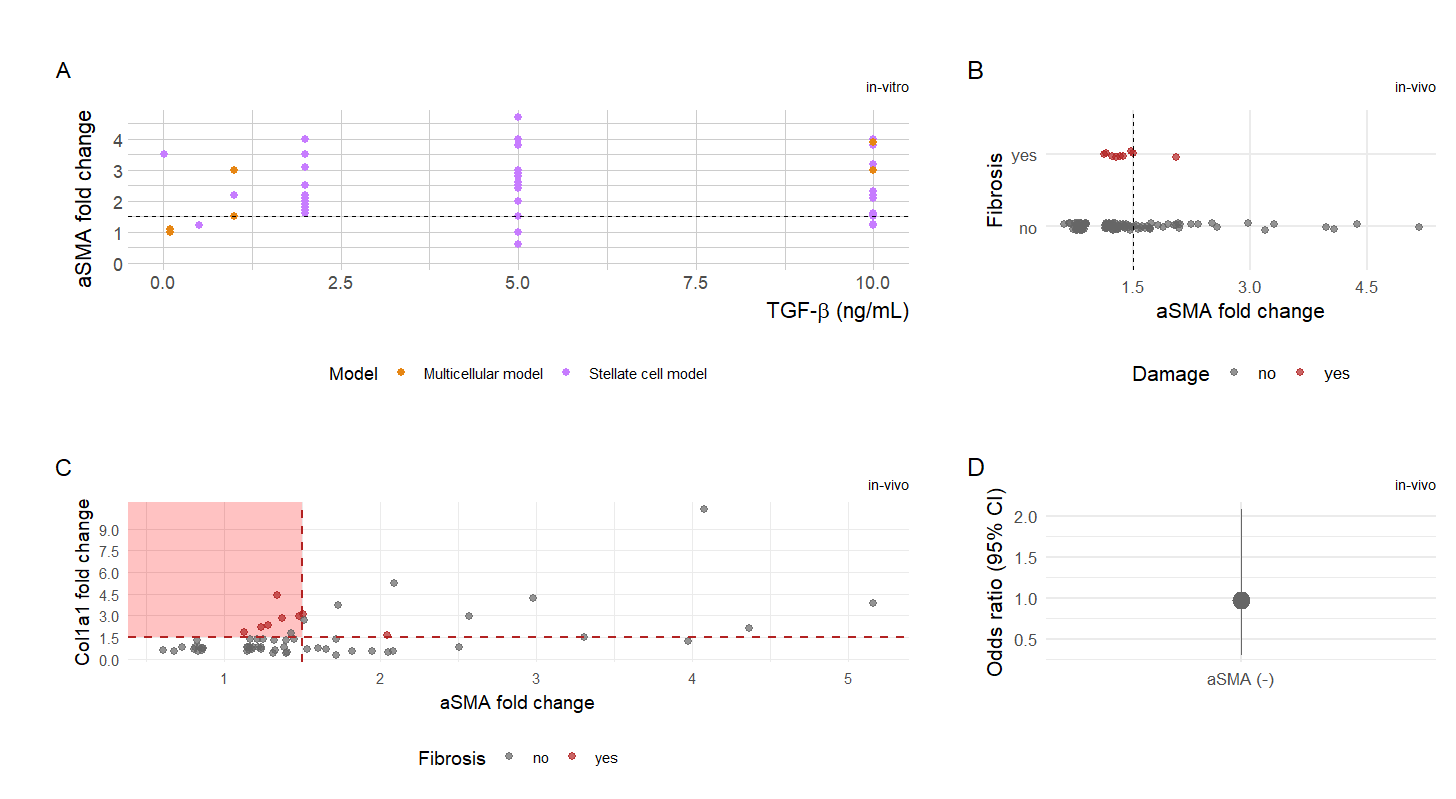
**Supplementary Figure S8:** Evaluating biologically relevant threshold for aSMA increase. A) Fold change in aSMA expression as a function of levels of TGF-β, used to activate stellate cells in vitro, in mono and multicellular in vitro models. One data point corresponds to a value reported in a given study for a given concentration. B) Fold change values aSMA from TG-Gates in vivo data plotted against damage status (Fibrosis), with points for individual observations. Points are colored by damage presence (grey = no, red = yes). C) Scatter plot of α SMA and COL1A1 fold changes in vivo, with points colored by fibrosis status. Grey points represent samples without fibrosis, and red points represent samples with fibrosis. Notice that most fibrotic cases are situated in second quadrant (in pink). D) Odds ratio with 95% confidence intervals from logistic regression models for aSMA fold change in vivo. Asterisks indicate statistical significance (p < 0.05) in neither (-), one (*) or both (**) tests (see Supplementary Table S4). Dashed lines in A, B, and C indicate the 1.5 fold change threshold.

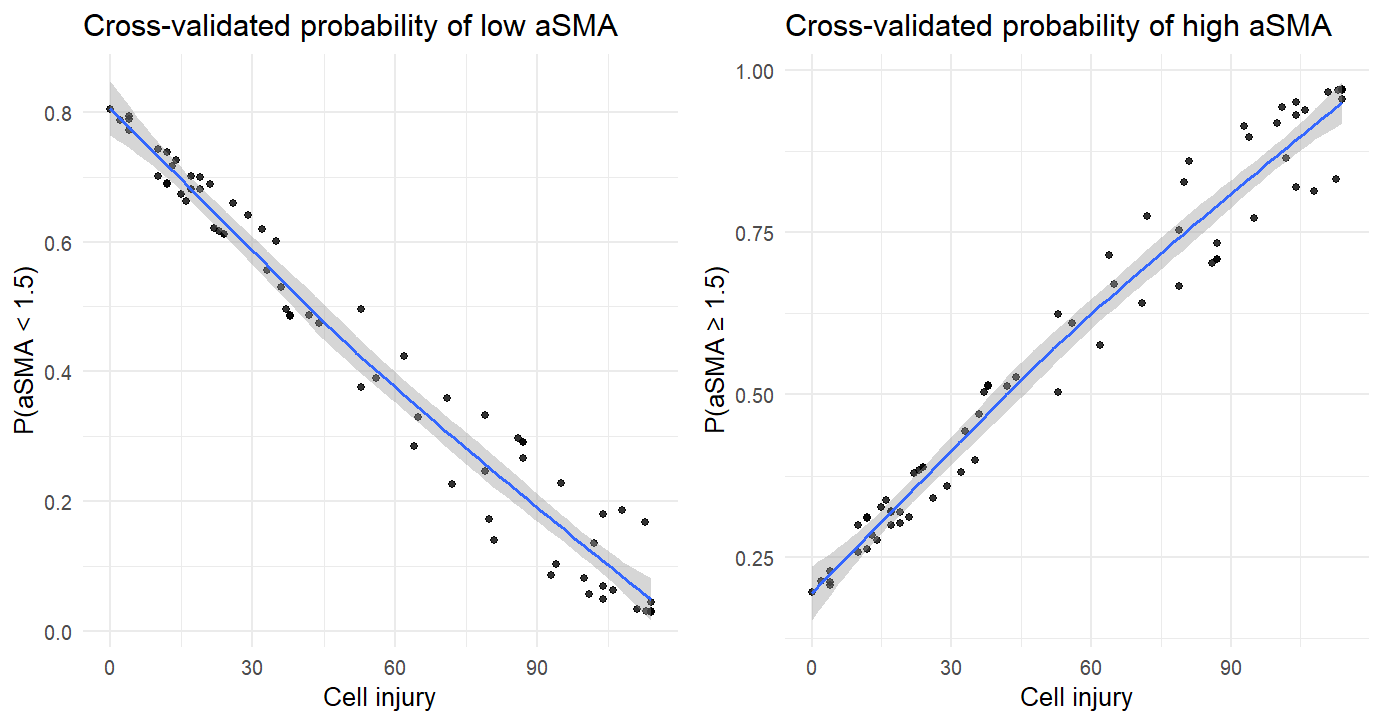

**Supplementary Figure S9: Conditional probabilities of aSMA levels given cell injury estimated from the Bayesian network model.** A) aSMA < 1.5 (lower HSC activation/liver fibrosis risk) and B) CSMAaOL1A1 ≥ 1.5 (higher HSC activation/liver fibrosis risk). Probabilities were obtained using 5‑fold cross‑validation. Points represent individual cross‑validated predictions, blue curves show LOESS smoothing, and grey shading indicates the 95% confidence interval across folds.

**Supplementary Table S1:** Summary statistics of aSMA, Col1a1, and ATP percentage in the cleaned dataset without outliers.Pctl X, Percentile X, SD, standard deviation.

| Variable | N | Mean | SD | Min | Pctl. 25 | Pctl. 75 | Max |
| --- | --- | --- | --- | --- | --- | --- | --- |
| ATP_perc | 69 | 59 | 39 | 0 | 21 | 96 | 114 |
| aSMA | 50 | 1.8 | 1 | 0.6 | 1 | 2.2 | 5 |
| Col1a1 | 57 | 2.5 | 2.4 | 0 | 0.9 | 3.1 | 12 |

**Supplementary Table S2:** Bootstrap estimates of Bayesian regression parameters with means and 95 % credible intervals computed from 1000 bootstrap resamples.

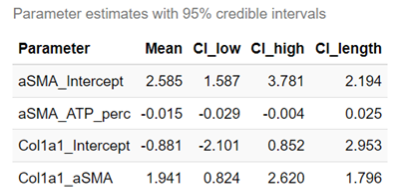

**Supplementary Table S3**: Fibrosis counts by marker significance (p < 0.1), NA = Not Available

| Significant marker | Fibrosis: yes | Fibrosis: no | Fibrosis: NA |
| --- | --- | --- | --- |
| COL1a1 | 14 | 133 | 904 |
| aSMA | 9 | 110 | 651 |
| Both | 8 | 53 | 219 |

**Supplementary Table S4:** Combined logistic regression and Wilcoxon results for (A) significant aSMA, (B) significant COL1A1 fold change from TG-GATES in vivo dataset

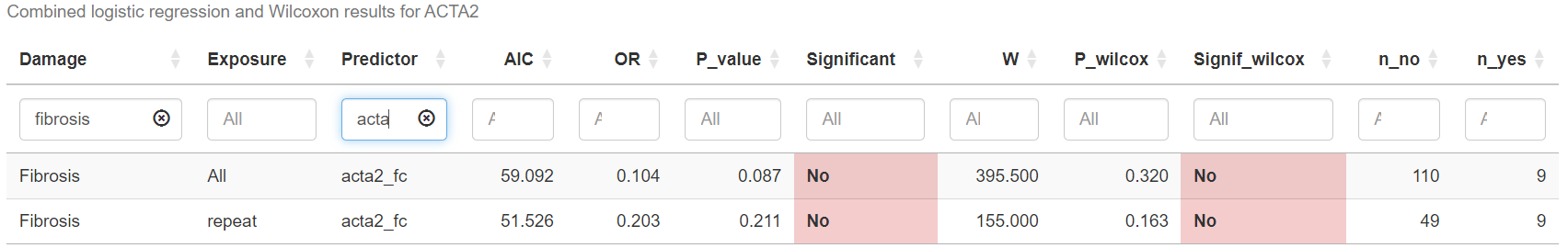

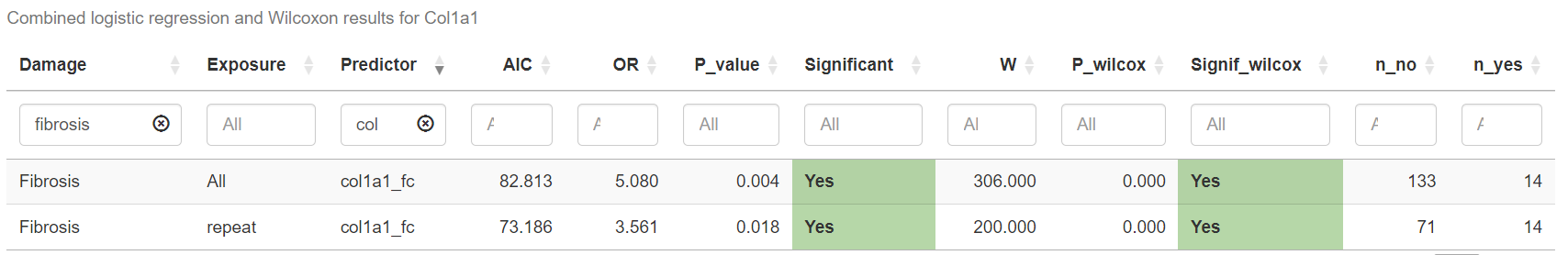
